# Signaling ligand heterogeneities in the peduncle complex of the cephalopod mollusc *Octopus bimaculoides*

**DOI:** 10.1101/2023.11.27.568875

**Authors:** Z Yan Wang, Clifton W Ragsdale

## Abstract

The octopus peduncle complex is an agglomeration of neural structures with remarkably diverse functional roles. The complex rests on the optic tract, between the optic lobe and the central brain, and comprises the peduncle lobe proper, the olfactory lobe, and the optic gland. The peduncle lobe regulates visuomotor behaviors, the optic glands control sexual maturation and maternal death, and the olfactory lobe is thought to receive input from the olfactory organ. Recent transcriptomic and metabolomic studies have identified candidate peptide and steroid ligands in the *Octopus bimaculoides* optic gland. With gene expression for these ligands and their biosynthetic enzymes, we show that optic gland neurochemistry extends beyond the traditional optic gland secretory tissue and into lobular territories. A key finding is that the classically defined olfactory lobe is itself a heterogenous territory and includes steroidogenic territories that overlap with cells expressing molluscan neuropeptides and the synthetic enzyme dopamine beta-hydroxylase.

## Introduction

Anatomical and physiological study of the *Octopus vulgaris* brain has attributed separate functions to each of the three peduncle complex territories, also called the supraoptic lobes [Messenger, 1967; Young, 1971]. The peduncle lobe is a visuomotor center that receives afferent connections from the ipsilateral optic lobe and basal lobes. All efferent connections from the peduncle lobe target motor centers [Young, 1971]. The olfactory lobe is intimately associated with the peduncle lobe: a continuous tract of neuropil joins the two lobes, making clear differentiation between the two difficult [Messenger, 1967]. Though “olfactory” in name, the function of the lobe is unclear and there is conflicting evidence as to whether this lobe processes olfactory information [Messenger, 1967; Messenger, and Young, 1997; Wells, 1978]. Indeed, the olfactory lobe in soft-bodied cephalopods appears to lack the glomerular organization that underlies olfactory processing in other taxa.

The third component of the peduncle complex is the optic gland. The spherical shape and yellow-orange color of the glands make them visually distinct from the two lobes. Functionally analogous to the vertebrate pituitary gland, the optic gland’s secretions control sexual maturation and reproductive behaviors [Wells, and Wells, 1959; Wells, and Wells, 1969; Wells, and Wells, 1975]. Removal of the optic glands in brooding female octopuses leads to a reversal of typical maternal behaviors and extension of lifespan [Wodinsky, 1977]. Recently, the optic gland secretions have been identified by transcriptomic and metabolomic analyses, revealing a rich interplay of molecular signals secreted across the lifespan of the octopus [Wang, and Ragsdale, 2018; Wang et al., 2022].

Despite the proposed multifunctionality of the octopus peduncle complex, little has been published on the internal anatomy of the peduncle complex since Young and Messenger’s work in the mid-20^th^ century. To better understand the peduncle complex, we assessed the supraoptic lobes of *Octopus bimaculoides* with modern molecular and bioinformatic techniques. First, we analyzed the peduncle complex of an *O. vulgaris* reference brain with that of our study species to establish species-specific understanding of the supraoptic lobe anatomy. We then assessed the *O. bimaculoides* peduncle complex for signaling molecule gene expression with *in situ* hybridization of known optic gland mRNAs [Di Cristo et al., 2003; Wang, and Ragsdale, 2018; Minakata et al., 2009; Minakata, 2010]. Our study reveals previously undescribed neurochemical heterogeneity in the peduncle complex of the adult female *O. bimaculoides*. Importantly, we found that labeling for key steroidogenic enzymes extends beyond that of the optic glands to include much of the olfactory lobe, a complex neuronal structure in which major peptidergic and catecholaminergic signals reside.

## Materials and Methods

### Animals

Wild-caught unmated female California two-spot octopuses (*O. bimaculoides*, n = 17) were purchased from Aquatic Research Consultants (Catalina Island, CA) and shipped overnight to Chicago, IL. Animals were individually housed in artificial seawater (Tropic Marin) in 20- or 50-gallon aquaria, and offered a diet of live fiddler shrimp, cherrystone clams, and grass shrimp. Water temperature was maintained between 17-21°C and water quality was checked daily. Ambient room temperature was 21-23°C. The animal room was kept on a 12:12 hour light/dark cycle.

### Perfusion, Tissue Processing, and Sectioning

Adult octopuses were deeply anesthetized in 5% ethanol/artificial sea water. The white body, a hematopoietic organ, was accessed by removing the top layers of skin on the head between the eyes. Animals were trans-orbitally perfused with freshly thawed 4% paraformaldehyde (PFA) in phosphate-buffered saline (PBS) delivered via a peristaltic pump (Cole Parmer Masterflex) through a 21½ gauge needle (Becton Dickinson). Left and right white bodies were targeted alternately and iteratively throughout the procedure. Following perfusion, the animal was decerebrated. The head was post-fixed overnight at 4°C in 4% PFA/PBS.

The next day, the central brain, together with the optic lobes, was dissected from the head mass. Brains were cryogenically protected in 30% sucrose/10% formalin in PBS at 4°C overnight, rinsed in 30% sucrose/PBS, and embedded in 30% sucrose/gelatin. Embedded brains were sucrose-sunk overnight in sucrose/formalin in PBS at 4°C, hemisected along the anterior-posterior axis, and stored at −80°C. The mantles of adult octopuses were dissected after tissue harvest to confirm sex and reproductive status. Only females with mature ovaries, ovarian follicles, and no evidence of fertilized eggs were used for subsequent analyses.

Tissue was sectioned coronally along the anterior/posterior axis on a freezing sledge microtome (Leica SM2000R) at a thickness of 24 microns, collected in diethyl pyrocarbonate (DEPC)-treated PBS, mounted onto Superfrost Plus slides (Fisher), dried overnight at room temperature, and frozen at −80°C until use.

### PCR isolation of genes

PCR primers for genes identified in the octopus genome or transcriptome were designed with MacVector software (version 12.6.0) (Table 1, Supplementary Materials). PCR reactions were conducted using a RoboCycler Gradient 40 (Stratagene). Reaction solutions were incubated at 94°C for 5 minutes before undergoing 35-40 rounds of amplification cycles: 94°C for 1 min, 52-55°C for 1 min, and 72°C for 1.25 minutes. The final elongation step was at 72°C for 10 minutes followed by storage at 9°C. PCR reactions were studied by gel electrophoresis to confirm that the sizes of products were as expected.

**Table 1.**
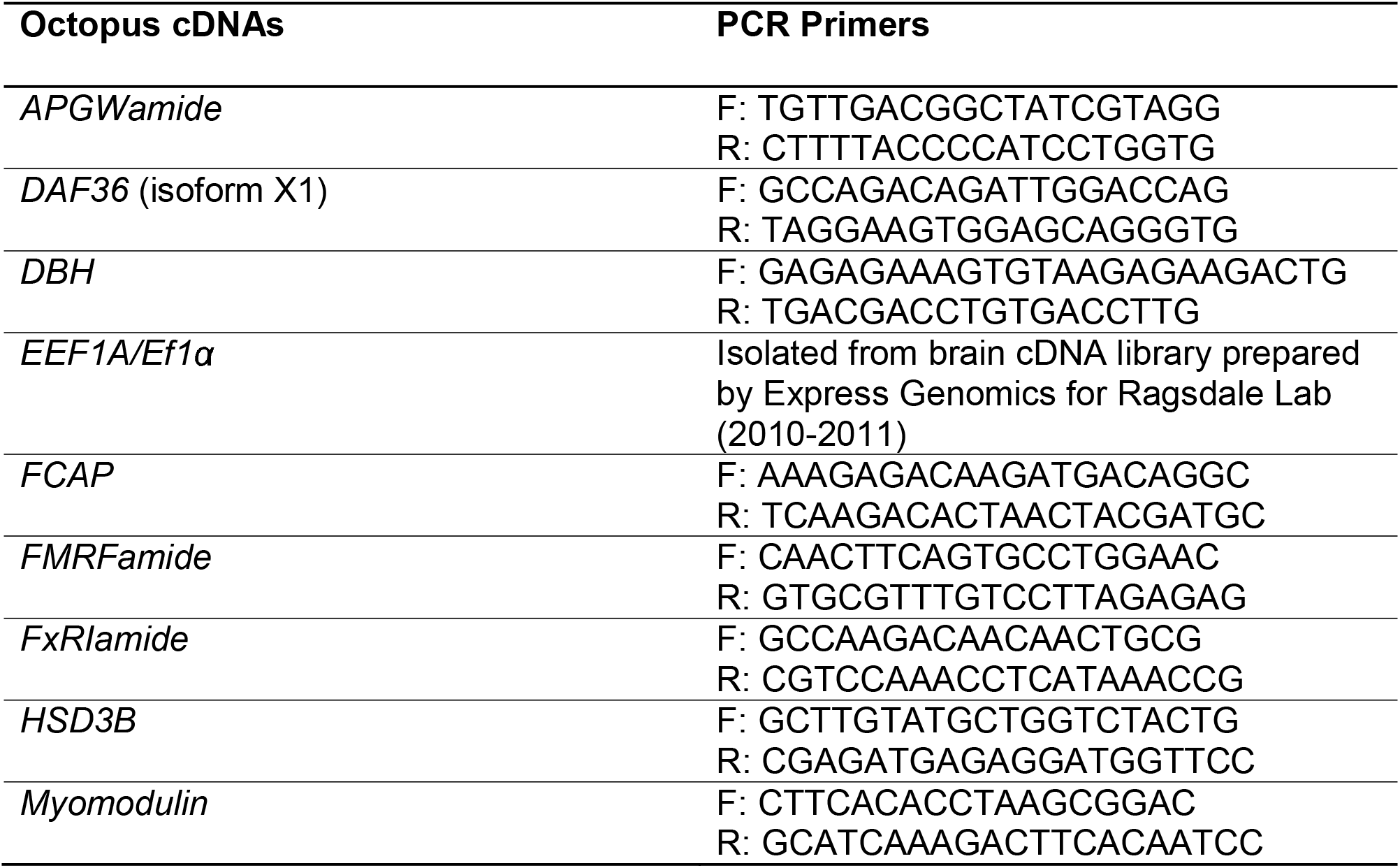
Molecular markers for analysis of octopus peduncle complex. F: forward PCR primer; R: reverse PCR primer.

### Riboprobe preparation

PCR reactions were ligated to pGEM T-Easy plasmid (Promega). Closed inserts were Sanger sequenced by the UChicago Comprehensive Cancer Center DNA Sequencing Facility. Plasmids were linearized by SacII or SpeI (NEB) restriction enzyme digestion to generate an antisense template. Following phenol-chloroform extraction of the template, antisense digoxigenin (DIG)-labeled riboprobes (Roche) were transcribed with SP6 or T7 RNA polymerase (NEB). After transcription, residual template was digested with RNase-free DNase I (Roche) at 37°C for 45 minutes. Riboprobes were ethanol-precipitated twice and stored in 100ul of 100% formamide at −20°C until use.

### *In situ* hybridization

Slides of sectioned tissue were equilibrated to room temperature and post-fixed for 15 minutes in 4% PFA/PBS, washed 3x five minutes in DEPC-PBS, and incubated at 37°C for 30 minutes in proteinase K solution (Roche Diagnostics; 1.5 μl stock proteinase K in 20ml of 100 mM Tris-HCl [pH 8.0], 50 mM EDTA [pH 8.0]). After proteinase K digestion, slides were post-fixed for 15 minutes in 4% PFA/PBS, washed 3x five minutes in DEPC-PBS, and stored in hybridization solution (50% formamide, 5x SSC, 1% SDS, 200 μg/ml heparin, 125 mg/ml yeast RNA, 0.5mg/ml acetylated BSA) at −20°C.

For the hybridization histochemistry, tissue was allowed to acclimate for at least one hour at 72°C in hybridization solution. Slides were transferred to mailers with 1-2 mg antisense riboprobe in 15 mL hybridization solution and incubated overnight at 72°C. The next day, slides were rinsed once in preheated Solution X (50% formamide, 5x SSC, 1% SDS) and washed 3x 45 minutes in Solution X at 72°C. Slides were next rinsed 3x 15 minutes in room temperature TBST (Tris-buffered saline [pH 8.5] with 1% Tween 20) and blocked for two hours in 20% DIG buffer (Roche) in TBST.

Incorporated DIG was detected with anti-DIG Fab fragments conjugated to alkaline phosphatase (AP, Roche). This conjugate was preadsorbed with octopus embryo powder in 5% DIG buffer in TBST for at least one hour. Slides were then incubated on a rocker overnight at 4°C in preadsorbed antibody diluted to a final concentration of 1:5000 in 5% DIG buffer in TBST.

The next day, slides were washed 3x 15 minutes then 2x one hour in TBST, followed by 3x 10 minutes in freshly prepared NTMT (100 mM Tris-HCl [pH 9.5], 100 mM NaCl, 50 mM MgCl2, 1% Tween 20). For the AP color reaction, slides were incubated in nitro blue tetrazolium (NBT, Denville Scientific) and 5-bromo-4-chloro-3-indolyl phosphate (BCIP, Denville Scientific) in NTMT. Color development proceeded at room temperature and was monitored for 1 hour to 5 days. Reactions were stopped in Stop TE (10 mM EDTA, 10mM Tris pH 7.5) and washed in TBST overnight. Slides were next fixed overnight at 4°C, dehydrated through an ethanol series, submerged in Histoclear (National Diagnostics), and coverslipped with Eukitt (Sigma-Aldrich). In our control experiments, digoxigenin-labeled RNA was omitted from the hybridization mixture. No signal was detected in the absence of digoxigenin-labeled riboprobes (not shown).

### Comparison of Octopus vulgaris and Octopus bimaculoides brains

To study the *Octopus vulgaris* central brain and optic lobes, we analyzed specimens from J.Z. Young’s collections at the Smithsonian National Museum of Natural History (USNM: 1593113; Accession number: 420332). Along with age and sex of the animals, the histological preparations of the tissue were not identified.

To describe the microanatomy of the *O. bimaculoides* brain, we performed *in situ* hybridization of octopus elongation factor 1-alpha (*EEF1A/EF1-alpha*), a broadly expressed molecular marker for animal cell bodies [Uetsuki et al., 1989]. We also stained brains with hematoxylin and eosin (H&E) to visualize nuclei and cytoplasm. Slides of sectioned tissue were equilibrated to room temperature and incubated overnight at 72°C in DEPC-PBS to remove gelatin. Following incubation, slides were rinsed twice in DEPC-PBS and once in DEPC-water. H&E staining was performed with slight modifications to manufacturer’s instructions (ScyTek). Briefly, slides were submerged in hematoxylin for 3.5 minutes, transferred to bluing reagent for 30 seconds, and then to eosin solution for 1 minute. Following dehydration, slides were coverslipped, as described above.

### Microscopy and image acquisition

Microscope slides were scanned by the Integrated Light Microscopy Core at the University of Chicago using an Olympus VS200 Slideview Research Slidescanner or a CRi Pannoramic SCAN 40x Whole Slide Scanner. Digital images were analyzed in QuPath [Bankhead et al., 2017].

### Identification of neuropeptide precursors

We performed a search for preproneuropeptide sequences with a custom perl script designed by Felipe Aguilera [Thiel et al., 2018], focusing on the most highly expressed transcripts of the optic gland to optimize the possibility of detection by *in situ* hybridization. We queried our data for the common cleavage site: [KR]-[KR]-x(2,35)-[KR]-[KR]-x(2,35)-[KR]-[KR]-x(2,35)-[KR]-[KR]. Cleavage probabilities were determined by NeuroPred, with mollusc as the model predictor [Southey et al., 2006]. Putative cleavage sites with a probability of cleavage below 0.9 were eliminated. We checked sequences for the presence of a signal peptide with SignalP [Nielsen, 2017] and confirmed identity of putative neuropeptides using BLASTP against the *O. bimaculoides* genome and non-redundant sequences in NCBI [Altschul et al., 1990]. DiANNA 1.1 web server was used to predict disulfide bridge formation [Ferrè, and Clote, 2006].

## Results

### Peduncle complex of O. vulgaris and O. bimaculoides

19th and 20th century studies of the *Octopus vulgaris* and *Eledone cirrhosa* central nervous systems divided the peduncle complex into the peduncle lobe, olfactory lobe, and optic gland [delle Chiaje, 1828; Young, 1971]. In recent years, the availability of new molecular methods has made *Octopus bimaculoides* the most suitable study species for neuroanatomical research [Albertin et al., 2015; Shigeno, and Ragsdale, 2015; Winters et al., 2020; Songco-Casey et al., 2022; Kang et al., 2023; Allard et al., 2023]. *O. bimaculoides* is a much smaller organism than *O. vulgaris*, and 31-24 million years of evolution separates the two species [Hanlon, and Messenger, 2018; Albertin et al., 2022]. Therefore, we were motivated to investigate whether any variation in neural organization exists at the level of the peduncle complex. We compared J.Z. Young’s *O. vulgaris* central nervous system specimens to our preparations of *O. bimaculoides* brains (Figure 1). To facilitate comparisons across different species and sample preparations, here we refer to three broad regions of the peduncle complex: anterior, intermediate, and posterior.

**Figure 1.**
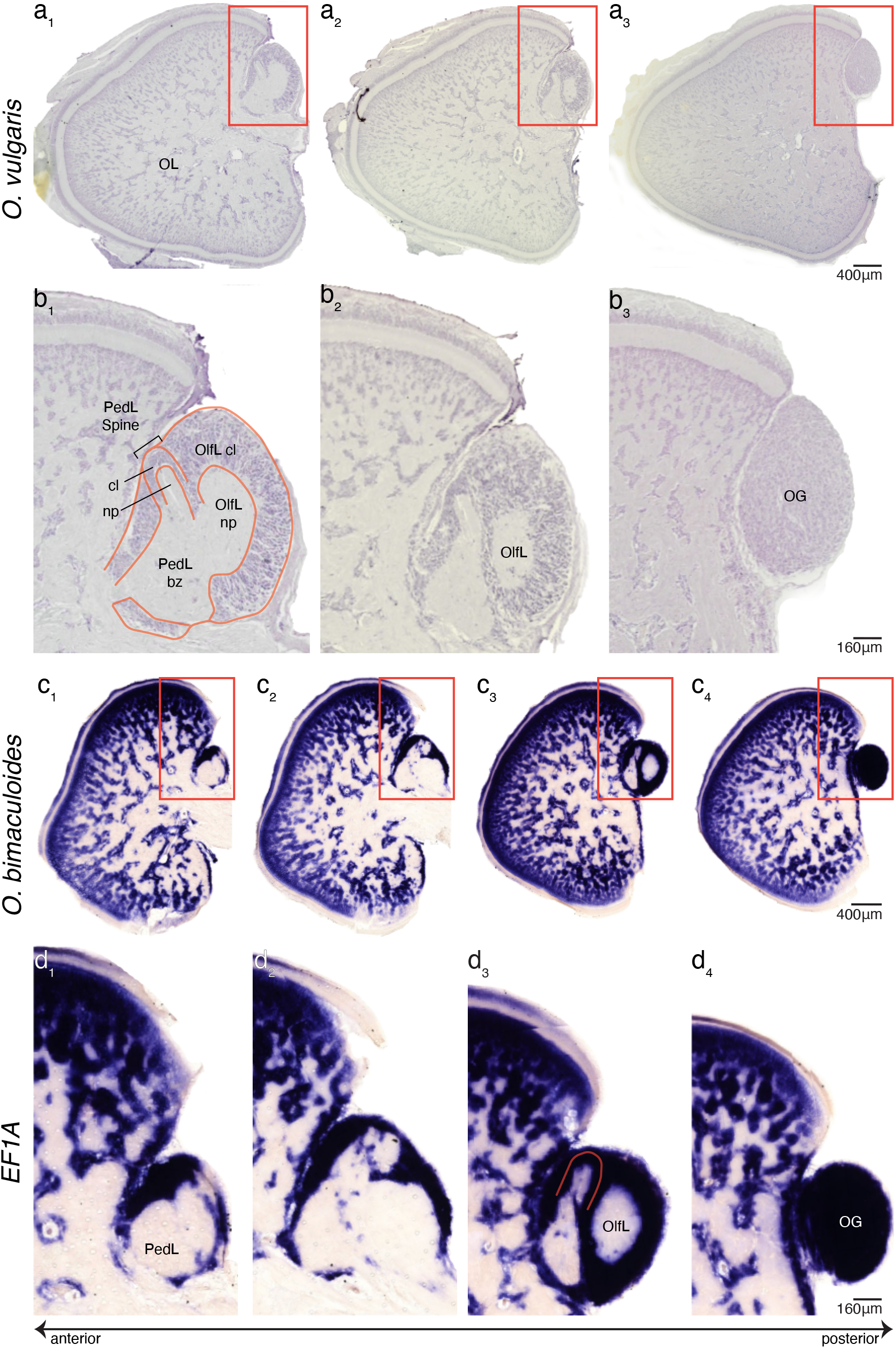
Anatomy of the octopus peduncle complex. Panels are arranged from left to right in the anterior to posterior direction. Regions outlined in a/c are depicted in higher resolution in b/d. **a-b.** *Octopus vulgaris* sections from J.Z. Young’s collection, unidentified neuronal stain. The spine of the peduncle lobe (**b_1_**) is clearly visible. **c-d.** *Octopus bimaculoides* sections, *EF1A in situ* hybridization. Candidate spine component of the peduncle lobe outlined in red (**d_3_**). PedL, peduncle lobe; OlfL, olfactory lobe; cl, cell layer; np, neuropil; bz, basal zone; OL, optic lobe; OG, optic gland.

Located anteriorly, the peduncle lobe comprises a spine and a basal zone. In *O. vulgaris*, the spine is a dorsal ridge of tightly-packed small cells whose fibers extend longitudinally through the majority of the lobe [Messenger, 1967; Young, 1971]. The spine is circumscribed ventrolaterally by the cell layer of the basal zone (also called the deep zone) and medially by the cell layer of the olfactory lobe (Figure 1b_1_), resulting in a well-defined, visually distinct neural territory. Within the lobular region of the *O. bimaculoides* peduncle complex, we also identify a lateral, peduncle territory and a medial, olfactory territory. The spine of the peduncle region appears to be greatly reduced in comparison to that of *O. vulgaris* (Figure 1c-d). A candidate spine component was identified near the posterior end of the peduncle lobe (Figure 1d_3_) but was not evident at more anterior levels (Figure 1d_1_-d_2_). This anatomy mirrors the H&E preparations of the *Octopus minor* central brain, which demonstrate a diminished spine that does not extend through the entire peduncle lobe [Jung et al., 2018]. Though *O. bimaculoides* is more distantly related to *O. minor* than to *O vulgaris*, *O. bimaculoides* and *O. minor* are much smaller in size when compared with *O. vulgaris* [Albertin et al., 2022; Jung et al., 2018]. The difference in peduncle spine anatomy is possibly explained by differences in body size and life histories across octopus species.

In both *O. vulgaris* and *O. bimaculoides*, the basal zone makes up the larger part of the peduncle lobe (Figure 1b_1_, Figure 1d_1_). Laterally, a cell layer separates the basal zone from the optic lobe. Medially, the neuropil of the basal zone is intimately associated with the olfactory lobe neuropil such that there is no clear demarcation between the two at anterior levels [Messenger, 1967; Young, 1971] (Figure 1b_1_). However, at more posterior levels, the cell layer of the olfactory lobe clearly separates the two lobes (Figure 1b_2_ and Figure 1d_3_). This organization is seen clearly in both *O. vulgaris* and *O. bimaculoides* (Figure 1).

At the most posterior end of the peduncle complex lies the optic gland [Messenger, 1967; Young, 1971] (Figure 1b_3_ and Figure 1d_4_). By Young’s neuronal stain and our *EF1A in situ* experiments, the optic gland appears as a spherical region of densely packed cells. A glandular territory, the optic gland grows as the octopus matures, and we observed variation in optic gland size across the animals studied [Budelmann et al., 1997].

Overall, by neuronal staining and *in situ* hybridization, the peduncle complex in both *O. vulgaris* and *O. bimaculoide*s includes a lobular area at the anterior and intermediate levels and a glandular region at the posterior end. We find the biggest difference in anatomy of in the anterior peduncle complex: the characteristic spine of the *O. vulgaris* peduncle complex is less pronounced in *O. bimaculoides*. However, clear differentiation between the putative optic gland region and lobular region is difficult to resolve through histological preparation alone. This in part motivated molecular and bioinformatic approaches to interrogating the anatomy of the territory.

### Steroidogenic enzyme gene expression extends beyond the optic gland

A role for the optic gland in cholesterol metabolism and steroid hormone production has been established recently with mass spectrometry and transcriptomic analysis [Wang, and Ragsdale, 2018; Wang et al., 2022]. These studies identified 3-beta-hydroxysteroid dehydrogenase (*HSD3B*) and an isoform of cholesterol 7-desaturase/Daf36 (*DAF36*), along with their steroid products such as 7-DHC, as being highly expressed in the optic glands. We followed up on these studies by performing *in situ* hybridization for *HSD3B* and *DAF36* on peduncle complex tissue sections.

We found that cells of the posterior peduncle complex show intense, homogenous labeling for both steroid biosynthetic enzymes in the optic gland (Figure 2). At the most posterior levels, this labeling accounts for the entirety of the cells in the peduncle complex. These results align with previously published descriptions of optic gland microanatomy and confirm recent transcriptomics data [Budelmann et al., 1997; Wang, and Ragsdale, 2018; Wang et al., 2022].

**Figure 2.**
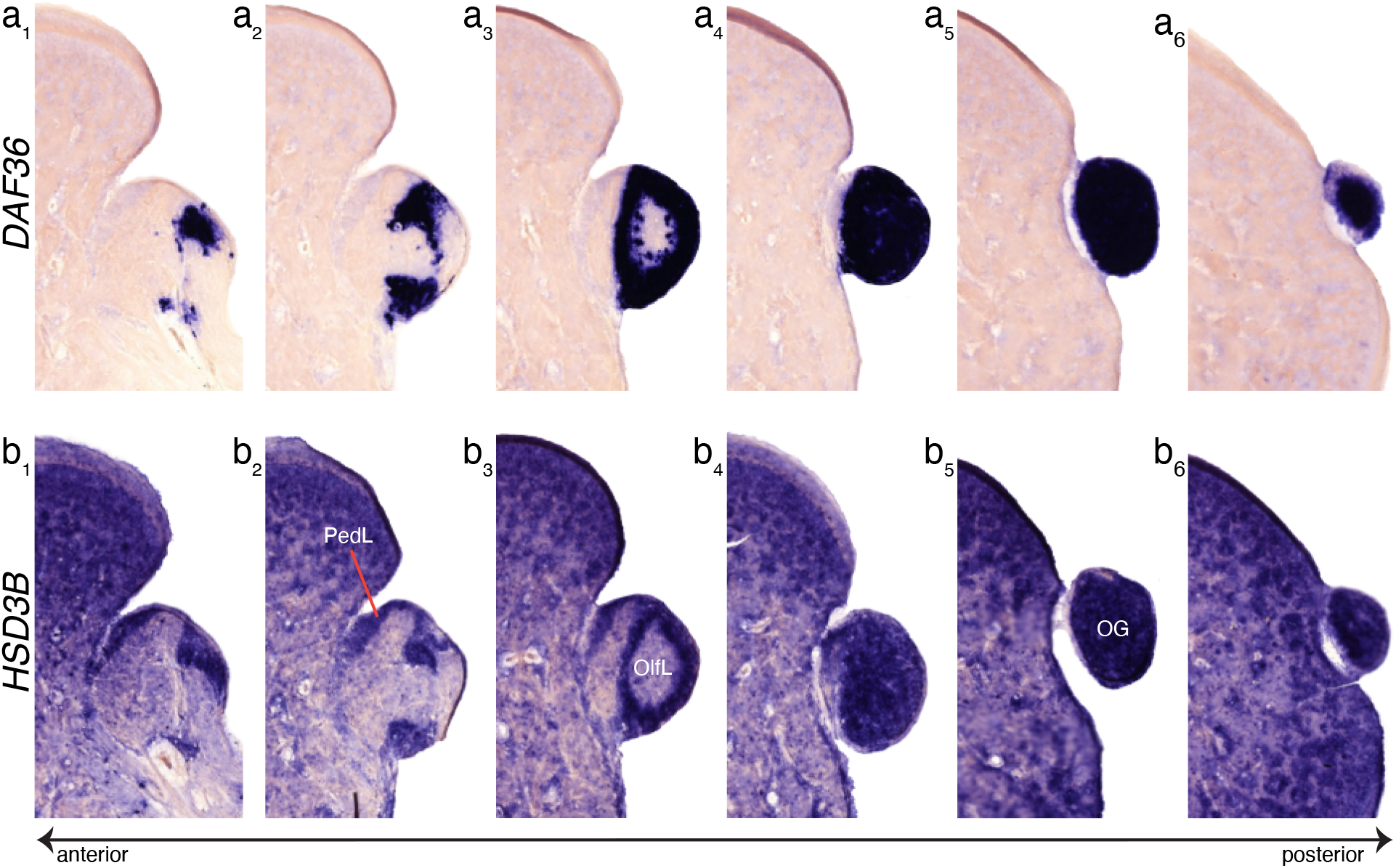
Steroidogenic enzyme labeling in the optic gland. *DAF36* (**a**) and *HSD3B* (**b**), two enzymes known to metabolize cholesterol into steroid hormones, show largely overlapping territories that extend from the olfactory lobe to the most posterior edge of the optic gland. *HSD3B* is expressed more broadly in the lateral peduncle lobe than *DAF36* (**b_2_**). Panels a/b are serially adjoining sections from the same animal. PedL, peduncle lobe; OlfL, olfactory lobe; OG, optic gland.

Surprisingly, we found that the robust steroidogenic signal extends anterior to the optic gland (Figure 2). *DAF36*-positive cells first appear at intermediate levels of the peduncle complex, corresponding to a part of the olfactory lobe (Figure 2a). *HSD3B* labeling shares this expression pattern, but also extends to the lateral cell layer of the peduncle lobe directly abutting the optic lobe (Figure 2b). Because our steroidogenic markers labeled only a portion of the cells at the intermediate peduncle complex, we sought to identify the molecular identity of the rest of the peduncle complex using ISH probes from other signaling families.

### Neuropeptide expression patterns in the peduncle complex

From optic gland transcriptomic data [Wang, and Ragsdale, 2018], candidate neuropeptide genes were analyzed with Neuropred and a protein motif searching program for the presence of signal sequences, cleavage sites, and short peptide repeats [Thiel et al., 2018] (Figure S1, Supplementary Materials). Identified homologs of FMRFamide, FxRIamide, myomodulin, LWamide, PRQFVamide, and APGWamide, and partial sequence for two amidated peptides were cloned with PCR and a subset were studied with ISH histochemistry (Figure 3).

**Figure 3.**
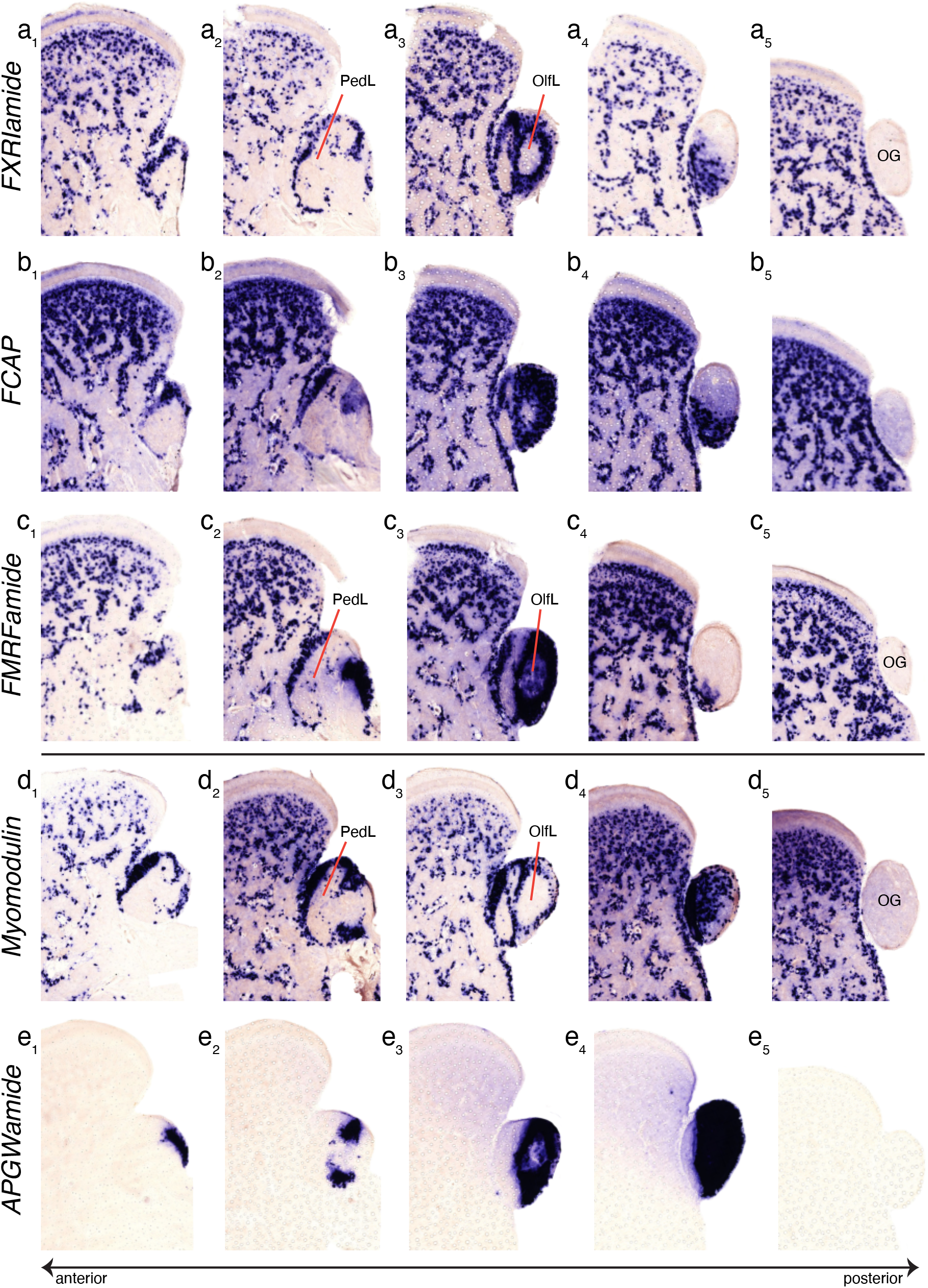
Neuropeptides define distinct compartments in the intermediate and anterior peduncle complex. No neuropeptide gene expression appears in the posterior peduncle complex, corresponding to the optic gland. Most neuropeptides we probed are also present in the cell bodies of the optic lobe neuropil and other regions of the central brain (see also Figure 5), but APGWamide is largely restricted to the olfactory lobe in the intermediate peduncle complex. Panels a/b and b/c represent serially adjoining sections from one animal. Panels d/e are serially adjoining sections from a second animal. PedL, peduncle lobe; OlfL, olfactory lobe; OG, optic gland.

The neuropeptide gene expression examined here formed two patterns: one shared by FXRIamide, FCAP, FMRFamide, and myomodulin, and one demonstrated by APGWamide (Figure 3). Gene expression for FXRIamide, FCAP, FMRFamide, and myomodulin was demonstrated throughout the lobes of the peduncle complex. At anterior levels, neuropeptide-positive cells were found at the lateral anterior cell layer of the peduncle lobe, with sparse labeling in the olfactory lobe cell layer (Figure 3a-d). Further posteriorly, labeling with the neuropeptide probes continues to the outer rind of the intermediate peduncle complex consistent with descriptions of the *O. vulgaris* olfactory lobe (Figure 3a_3_, Figure 3c_3_, Figure 3d_3_). At this level, these four markers also pick out scattered cells in the peduncle and olfactory neuropil (Figure 3a_2_, Figure 3c_2_, Figure 3d_2_). At the posterior edge of this neuropeptide territory, the four probes show strong but dappled labeling, suggesting that cells producing different neuropeptides may be interposed amongst each other (Figure 3a_4_-3d_4_). This heterogeneous neuropeptide region retreats ventrolaterally, disappearing completely at the level of the optic gland (Figure 3a_5_-3d_5_).

APGWamide, one of the most highly expressed transcripts in the female optic gland, is not expressed in the peduncle lobe, unlike the other four neuropeptides (Figure 3e_1_) [Wang, and Ragsdale, 2018]. Strikingly, at anterior levels, APGWamide labeling appears in a small zone of cells located dorsomedially in the olfactory lobe (Figure 3e_1_). Interestingly, while the other neuropeptide markers are expressed in the cells of the optic lobe neuropil (Figure 3a-d) and other regions of the central nervous system (Figure 4, Figure 5), APGWamide expression is restricted to the olfactory lobe (Figure 3e). This aligns with immunohistochemistry experiments in *O. vulgaris*, in which APGWamide immunopositive neurons are primarily found in the posterior olfactory lobe [Di Cristo et al., 2005]. APGWamide message was not detected at the most posterior end of the complex where the optic gland sits and where the steroidogenic markers were detected (Figure 3e_5_).

**Figure 4.**
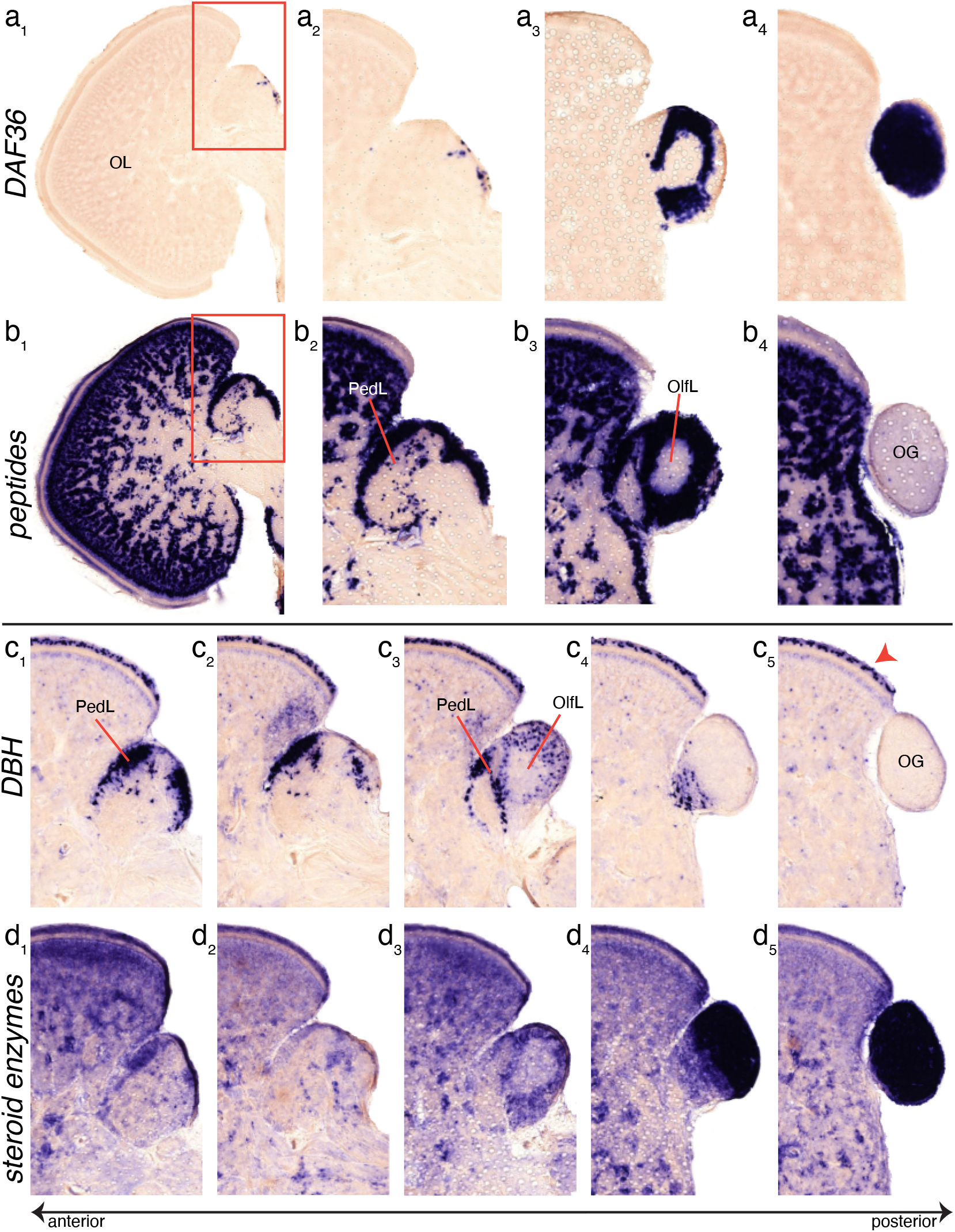
Interposed neuropeptide and steroid *in situ* hybridization. **a-b.** The intermediate level of the peduncle complex has both *DAF36*-positive and neuropeptidergic territories, with highest similarity in expression pattern at the olfactory lobe. Regions outlined in **a_1_/b_1_** are shown in higher resolution in panels **a_2_/b_2_**, respectively. “Peptides” includes RNA probes for FMRFamide, FxRIamide, FCAP, and myomodulin. **c-d.** Labeling for *DBH* and steroid enzymes (*HSD3B* and *DAF36*) overlaps in the olfactory lobe and lateral posterior peduncle lobe but is otherwise dissimilar. *DBH* expression is strongest in the lateral posterior peduncle lobe (**c_1_**) and the outer granular layer of the optic lobe (arrowhead in **c_5_**). Panels a/b are serially adjoining sections from the same animal. Panel c/d are serially adjoining sections from a second animal. PedL, peduncle lobe; OlfL, olfactory lobe; OG, optic gland.

**Figure 5.**
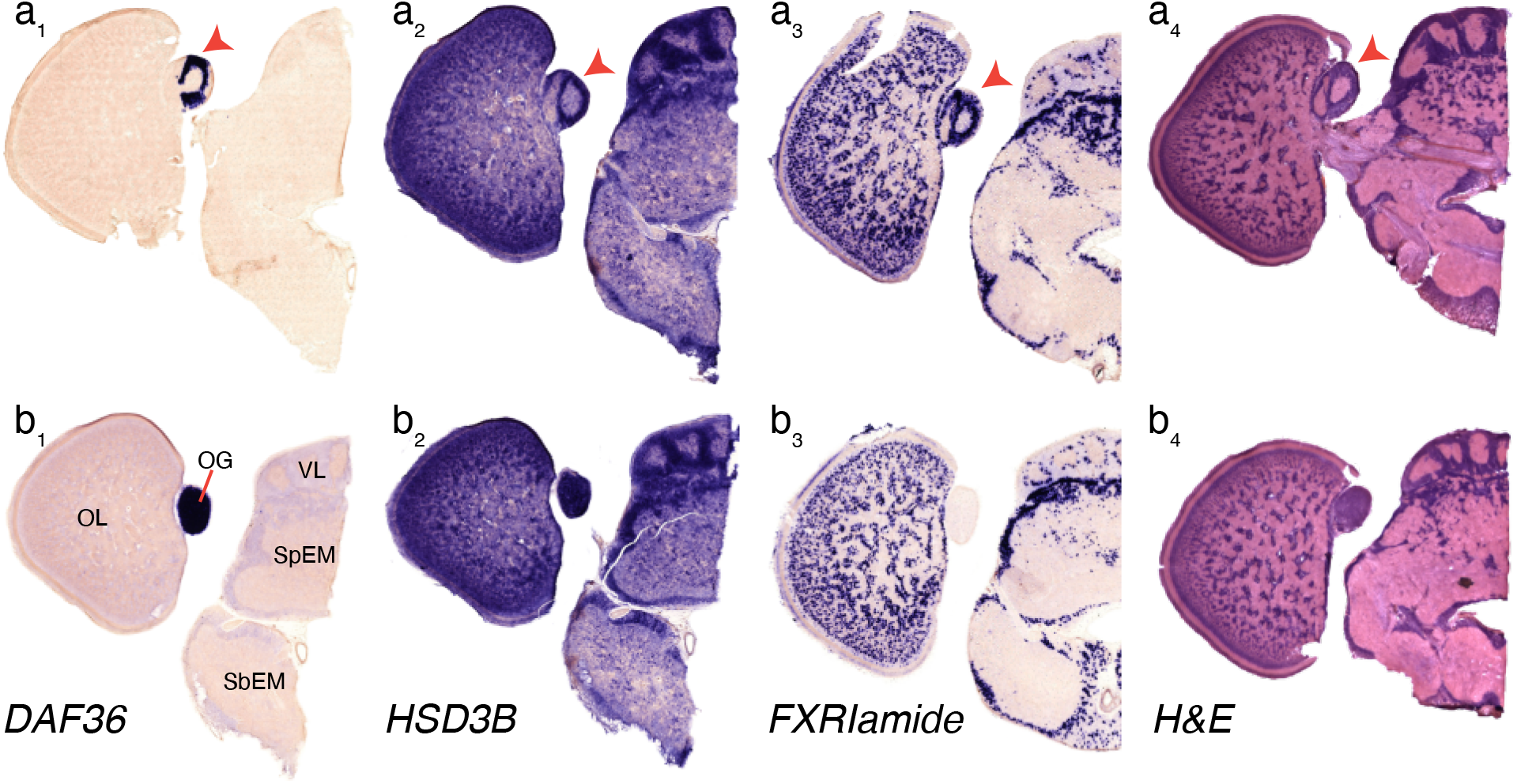
Neurochemical complexity in the octopus peduncle complex. The steroidogenic enzyme genes *DAF36* (**a_1_**, **b_1_**) and *HSD3B* (**a_2_**, **b_2_**) are expressed in the olfactory lobe (arrowheads) and throughout the optic gland. *DAF36* is restricted to the peduncle complex. *HSD3B* expression is more widespread in the nervous system, including the vertical lobes (**a_2_**, **b_2_**). *HSD3B* is also found in the neuropil, suggesting the possibility of glial cell expression. Neuropeptides, such as FXRIamide (**a_3_**, **b_3_**), are expressed throughout the nervous system and in the peduncle and olfactory lobes, but are completely missing in the optic gland (**b_3_**). Hematoxylin and eosin stain (**a_4_**, **b_4_**) shown for reference. OL, optic lobe; OG, optic gland; VL, vertical lobes; SpEM, supraesophageal mass; SbEM, subesophageal mass. Sections are representative slides from several animals and are not serially adjoining.

Overall, these results reveal a rich and varied population of neuropeptidergic cells in the intermediate level of the peduncle complex. Neuropeptides with broad distribution in the nervous system and diverse putative functions are also found more anteriorly in the peduncle lobe [Cropper et al., 1987; Wang et al., 1998; Jing, and Weiss, 2001; Sweedler et al., 2002; Veenstra, 2010]. However, APGWamide is specific to the olfactory lobe [Di Cristo et al., 2005]. These findings suggest a neurochemical correlate for the divergent functions of the olfactory and peduncle lobes.

### Collective distributions of steroidogenic, neuropeptidergic, and catecholaminergic message in the peduncle complex

We directly tested whether the steroid and neuropeptide territories of the peduncle complex overlap with each other by performing ISH experiments with alternating sections probed for *DAF36* and for all the studied peptides (Figure 4a-b). We found that the two signaling classes marked partially separate territories: a neuropeptidergic-rich area in the anterior and intermediate optic gland that gives way to a territory at the posterior peduncle uniformly labeled by cholesterol-metabolizing enzyme probes (Figure 4a-b). Any cellular overlap that exists between peptidergic and steroidogenic enzyme expression would lie in the cell layer of the olfactory lobe.

We explored whether the steroidogenic cells shared expression territory with that of dopamine beta-hydroxylase (*DBH*), which had been previously identified as differentially expressed in the optic gland transcriptomes [Wang, and Ragsdale, 2018]. DBH is the rate-limiting enzyme in the production of monoamine neuromodulators, including norepinephrine in vertebrates and octopamine in invertebrates [Juorio, and Molinoff, 1971]. Our results with serially neighboring sections show that the most robust *DBH* labeling is found at the dorsal lateral cell layer of the peduncle lobe and medial cell layer of the olfactory lobe (Figure 4c_1_). At the intermediate level of the peduncle complex, there is robust labeling for *DBH* in the peduncle lobe cell layer and moderate labeling in the olfactory lobe cell layer (Figure 4c_3_). In comparison with the neuropeptide labeling, only scattered cells in the perimeter of the olfactory lobe are *DBH*-positive (Figure 4c_3_). *DBH* expression has some overlap with that of the cholesterol metabolizing enzymes at the anterior level of the peduncle complex, where *HSD3B* but not *DAF36* is expressed (Figure 4c_1_, 4d_1_). We also detect *DBH* in the outer granular layer of the optic lobe (Figure 4c_5_), and in a few cells of the optic lobe neuropil. Our findings confirm transcriptomic results localizing *DBH* in the octopus nervous system and suggest that DBH is broadly important for signaling in multiple neurochemical regions of the peduncle complex [Juorio, and Molinoff, 1971; Songco-Casey et al., 2022].

## Discussion

Here, we report findings from ISH and histochemical experiments of the peduncle complex that augment recent transcriptomic and metabolomic studies of the octopus neuroendocrine system. We confirm the presence of multiple signaling systems in the three peduncle complex territories. In the unmated female optic gland transcriptome, we previously detected both steroidogenic enzyme message as well as message for neuropeptides [Wang, and Ragsdale, 2018]. Our neuropeptide ISH experiments here confirm the presence of neuropeptides in the peduncle and olfactory lobes, but cannot distinguish the cellular layer of the posterior olfactory lobe from the cells of the optic gland (Figure 3). It is possible that the bulk tissue dissections for transcriptomics included nearby lobular tissue. This problem is accentuated in *O. bimaculoides* because the transition between the olfactory lobe and the optic gland is not evident (Figure 1). The boundary between lobular and glandular regions of the peduncle complex is further obscured by our finding that steroid enzymes are not clearly restricted to the optic gland (Figure 2). At present, there does not exist a molecular marker specifically coextensive with either the optic gland or olfactory lobe [Di Cristo et al., 2005]. Therefore, the extent, if any, to which neuropeptidergic cells constitute some small district within the optic gland remains an important open question.

Both ISH and H&E data show homogeneous distribution of steroidogenic enzyme positive cells in the optic gland, which matches previous anatomical descriptions of the region [Budelmann et al., 1997] (Figure 5). DAF36 and HSD3B are cholesterol-metabolizing enzymes that appear responsible for the steroid changes in mated female *O. bimaculoides* [Wang, and Ragsdale, 2018; Wang et al., 2022]. The strong consensus in labeling of the two probes suggests that intracellular steroid biogenesis occurs in the posterior peduncle complex preceding release or downstream processing. Strikingly, the glandular region marked by steroidogenic enzymes extends beyond the boundaries of the optic gland proper into the olfactory lobe, where they interface seamlessly with *DBH*- and neuropeptide-positive cells (Figure 2, 5).

Of the neuropeptides we examined, APGWamide mRNA is unique for its restricted distribution in the olfactory lobe. In gastropod molluscs, APGWamide modulates copulatory and other reproductive behaviors [de Lange, and van Minnen, 1998; Oberdörster et al., 2005]. Our ISH results align closely with a previous study in *O. vulgaris* that localized abundant APGWamide immunopositive cells in the olfactory lobe [Di Cristo et al., 2005]. The confined region of APGWamide labeling in the peduncle complex overlaps with positive labeling for steroidogenic enzyme message, strongly suggesting that reproduction is a major function of the olfactory lobe.

The olfactory lobe was named such because it receives nerve fibers from the so-called olfactory organ or pit, which has large, ciliated cells in the outer epithelial layer reminiscent of a chemoreceptive organ [Woodhams, and Messenger, 1974]. However, in octopus, a putative olfactory function of the lobe has never been confirmed with behavior studies. Indeed, as many researchers have noted, little evidence supports that either the octopus olfactory organ or lobe are involved in chemosensation [Messenger, 1966; Young, 1971; Wells, 1978; but in squid, see Gilly and Lucero, 1992]. The name is retained, although as early as 1967, researchers have proposed putative neurosecretory activity in olfactory lobe cells based on their histological characteristics [Bonichon, 1967]. Our findings suggest that the olfactory lobe participates in steroidal and neuropeptidergic signaling, providing a molecular basis for the reproductive functions of the olfactory lobe. The other neuropeptides examined here are expressed throughout the cell layer of the peduncle lobe and the other central nervous system lobes (Figure 3, Figure 5). Some structures marked by these neuropeptides are likely involved in feeding, as they are in gastropods [Cropper et al., 1987; Wang et al., 1998; Jing, and Weiss, 2001; Sweedler et al., 2002; Veenstra, 2010], but their broad distributions point to a more general role in neuronal transmission.

## Acknowledgements

We thank Dr. Chuck Winkler of Aquatic Research Consultants for providing us with octopuses, Dr. Michael Vecchione and Kathryn Ahlfield of the National Museum of Natural History for their stewardship of Young’s archives, Dr. Christine Labno of the UChicago Integrated Light Microscopy Core for assistance with slide scanning, and Drs. Andreas Hejnol and Felipe Aguilera for sharing code ahead of publication.

## Statement of Ethics

All research was performed in compliance with the EU Directive 2010/63/EU guidelines on cephalopod use [Fiorito et al., 2014; Fiorito et al., 2015; Lopes et al., 2017] and the UChicago Animal Resources Center and Institutional Animal Care and Use Committee.

## Conflict of Interests

The authors declare no conflicts of interest.

## Funding Sources

This research was supported by NSF grant IOS 1354898 to C.W.R.

## Author Contributions

Conceptualization: Z.Y.W, C.W.R.; Investigation: Z.Y.W.; Writing - Original Draft: Z.Y.W.; Writing – Review & Editing: Z.Y.W, C.W.R.; Funding Acquisition: C.W.R.

## Data Availability Statement

Data are available upon reasonable request to the authors.

---

**Figure S1. Prepropeptide sequences predicted from the optic gland transcriptomes.**

The putative signal peptides are boxed and highlighted in yellow, predicted cleavage sites are in red, and known diagnostic peptide sequences are in blue. In amidated peptides, glycine residues to be converted into C-terminus amides are in green.

## Notes

### Competing Interest Statement

The authors have declared no competing interest.

