## Supplemental Figure 1 for "Signaling ligand heterogeneities in the peduncle complex of the cephalopod mollusc *Octopus bimaculoides*"

a

**FMRFamide**

MSGLSPFSLLIFIFIYCLTVYAATAMSLAQACMESPSMCESISLSHLSSEEGLKSKRFLRPGRALSGDAFLRFGKNGLNL
PFEDKRFLRFGRTDRQFEDMLKEVLQRAENVDNRQKRSTDNIPQSSVQDDSSKITKRNADASDNGMDKRFMKFGKSGDLP
YVNEWSDKRFMRFGREPDKRFMRFGKSDDKRFMRFGRNPNELEDRLEEGKRFMRFGRGNEEEEKRFMRFGRDPDSKLMGY
GNSEEEKRFMRFGRMSDEADAQKRFMRFGRSVDDDKANRLKKSNDQLRTIRMGRSVDDKKVDRLKTSNDRLPVITMGRSV
GDKTVKKL*

b

**FxRIamide**

MAGLWRIVLLATITLMCVTSQWVIYVQAEEASSSDELVNSANEADPEMDSEDDLKRANTFLRIGKANGILRLARSPSSFL
RIGRRPPLHFVRIGKAPSSMFLRIGKRVDDSENDDLNEYDADKMSGRDTRASPSSFLRIGKSGNEVEDEIDDTDETVKRV
NAFLRIGRQNDPSSFLRIGKSLNNDDLSKDKRTNAFLRIGKIPASSFIRLGRGPFTEDNGINTRGFRGPTRGFLRIGKRA
AIPDGSHADYFSDLNVKSQ*

c

**FCAP**

MTTLPRHKIVLCRLVNREKELRRIMPTGYRSSKNGKILISEFIVLTKGSNHLSPSKKRKILSPHNEDNQFETIKMKLKGK
CNVSFIVFEEHDKERRSPNGCNEKRQLVDKGPIGKLKNDEFDAAYWNTDSKNMNKRLSDSVKDLPIGMHVLHRKRSFDSL
GGAFMPGGKRNFDSLGGAFMPGGKRNFDSLGGAFMPGGKRNFDSLGGAFMPGGKRNFDSLGGAFLPGGKRNFDSLGGAFM
PGGKRNFDSLGGAFMPGGKRNFDSLGGAFMPGGKRNFDSLGGAFMPGGKRNFDSLGGAFMPGGKRNFDSLGGAFMPGGKR
NFDSLGGAFMPGGKRTFDSLGGGILPIEGSIDLTSGCLGKRSCHSLSGRRLSAGSSADSSDDSDGKRSFDSLGGAFMPGG
KRTFDSLGGAFMPVGKKNFDSLGGAFMPGGKRTFDSLGGAFMPGGKRNFDSLGGAFMPGGKRNFDSLGGAFMPGGKRTFD
SLGGAFMPGGKRTFDSLGGAFMPGGKRNFDSLGGAFMPGGKRTFDSLGGAFMPGSKKNFDSLGGAFMPGGKRQSDSLGGG
KRQFDSLGGAFMPGGKRQFDSLGGAFMPGGKRQFDSLGGAFMPGGKRQFDSLGGAFLPNIKRQDDRPYLWQPFDSLGGAF
LPGGKRDSDSLSGAILTESAEKQGSGDSDLKN*

d

**Myomodulin**

MLFNITPIICALVLCVLFPRGNCNEDTNASQSKSKLNENNVAEQHLDKRSSGEYNPNDLKTIVAAILERQEQGQKQSSAL
RSMNELAGDQYNGDALSRVVQLLRRSMPSDFDDFERRLAPVPRLGRLKKRSVLSNPNKEVDYDEIEANDNNESADNILFP
RQITLPRYGKDEGSNAADAEQYLSSDCEVFDAYGTCVQYKIPGENVKRQVRMLRLGKRQMHPIKIDDESTGMDKRSEDFN
ADGNSGENKRAVSMLRLGRSFNAGGQSEENEIPVFDEAKRAVSMLRLGRSGPFLDKRALSMLRLGRSEFDQENTALPILY
PGQNGFENNKRAVSMLRLGRSMSAGDKRAVSMLRLGRSGFDTMKRAVSMLRLGRNSGYPSEKRAVSMLRLGRSGSDEDKR
AVSMLRLGRSGADIEDEKRAVSMLRLGRSGADNMKRAVSMLRLGRSGSDDMNTKRAVSMLRLGRSGNDNVGEDKRAVSML
RLGRSDNNANNKRAMAMLRLGRSNDTSSKET*

e

**APGWamide**

MMLTAIVGIRNYARFASAKIFWYAALILIAYSLTSTTNNAVAGSDSTTIELKTDVKRGKRSVDASQHSDQHHVHRYNGKQ
SQQQAHASISALAPIPSELDSYPNDFSSIDFNSYDAKPEEVFTGNQDGEVGQLFVIKQSSGNEETEETEGNDITEELSSK
ADLSNLNLATEAEKRAPGWGKRAPGWGKRAPGWGKRAPGWGKRAPVLRKREFEITTPELGDLDFQKRAPGWGKRDHFSLG
LN*

f

**Allatostatin/LWamide-like**

MTPSSMWMKVLFCSILLHLYVAQYTRSLETHSEDDSESDKRSIIDDDFKSLDPTYYPSGLDKRQQQYSKLGRNIRVIARG
MDPMMFGNLGKRMDPNMFGSLGKRFNTDGREKQDKKIDPYMFGSLGKRMDPAMFGSLGKRMDPMLYGSLGKRMSPDYINY
AGKRYDPVLFGGLGKRMDPMLFGGLGKKMDPNMFGTLGKRMDSMMFGHLGKRDVSGIKVGDFE

g

**IRELVamide/PRQFVamide-like**

VGKRGTEVEFPDKRIRELIGKRGGEIDYPDKRIRELIGKRGGEVEYPENFEKRIRELIGKRGTEIEFPEQVDKRIRELIG
KRGTEVEYPEHFDKRIRELVGKRYPENFDKRIRELVGKRGTEIDYPDHFDKRIRELVGKRGTEIEYPDHFDKRIRELVGK
RGNEVEYSDHFDKRIRELVGKRGTEVEFPDKRIRELIGKRGGEIDYPDKRIRELIGKRGGEVEYPENFEKRIRELIGKRG
TEIEFPEQVDKRIRELIGKRGTEVEYPEHFDKRIRELVGKRGNEAENPEHFDKRIRELVGKRSVPSNISKNVVKRNVEKL
EG

**Figure S1. Prepropeptide sequences predicted from the optic gland transcriptomes.** The putative signal peptides are boxed and highlighted in yellow, predicted cleavage sites are in red, and known diagnostic peptide sequences are in blue. In amidated peptides, glycine residues to be converted into C-terminus amides are in green.
