## Supplemental Materials for "Signaling ligand heterogeneities in the peduncle complex of the cephalopod mollusc *Octopus bimaculoides*"

### Supplementary Materials

**Details on octopus peptide sequences**

*FMRFamide:* FMRFamide was previously identified in the optic glands by immunohistochemistry [Di Cristo et al., 2003]. SignalP prediction software predicted a signal peptide 22 amino acids in length (Figure S1). The octopus FMRFamide sequence shows a single copy of the decapeptide ALSGDAFLRFamide, a single copy of FLRFamide, and 7 copies of FMRFamide. The octopus FMRFamide-related peptides bear sequence similarity to other FaRPs that have recently been identified in cuttlefish and other molluscs [López-Vera et al., 2008; Zhang et al., 2012; Ahn et al., 2017; Li et al., 2018].

*FxRIamide:* FxRIamide was initially identified by a bioinformatics screen of the whole-body transcriptome of the gray garden slug, *Deroceras reticulatum*, and resembles the FaRPs that have been reported in several lophotrochozoans [Ahn et al., 2017]. FxRI is similar in sequence to FMRFamide, suggesting it may play related roles in modulating synaptic activity. Octopus FxRIamide has a 28 amino acid predicted signal peptide sequence and 3 copies of the heptapeptide xSSFxRI, along with several closely related peptide sequences (Figure S1).

*Feeding circuit activating peptide (FCAP):* Originally identified in the buccal neurons of *Aplysia*, FCAP is an amidated neuropeptide thought to initiate rhythmic feeding motor programs and modulate food-related arousal [Sweedler et al., 2002]. *FCAP* encodes multiple copies of closely related undecapeptides, with the major form NFDSLGGAFMPamide appearing 16 times in the octopus sequence (Figure S1).

*Myomodulin:* We recovered an octopus homolog of myomodulin, which was previously unannotated in the genome (Figure S1). Myomodulin was identified in buccal motor neurons of *Aplysia*, where it was found to potentiate contraction of the accessory radula closer buccal muscles [Cropper et al., 1987]. We found a predicted signal sequence of 23 amino acids and 12 copies of AVSMLRLamide and two nearly matching variants. This peptide sequence is similar to *Aplysia* myomodulin, which has a sequence of PMSMLRLamide [Cropper et al., 1987].

*APGWamide:* APGWamide activity has been implicated in the control of male copulatory behavior in gastropod molluscs [de Lange and van Minnen, 1998]. In *O. vulgaris*, APGWamide-immunoreactive fibers have been found in lobes of the central brain, the glandular cells of the oviducal gland, and neurons of the posterior olfactory lobe, suggesting a shared reproductive function in cephalopod molluscs [Di Cristo et al., 2005; Minakata, 2010]. The octopus homolog contains several copies of APGW at the 3’ terminus (Figure S1).

*Other neuropeptides*: We also found partial sequences for two amidated peptides (Figure S1). Allatostatin/LWamide was previously reported in octopuses [Wang and Ragsdale, 2018] and is known to modulate feeding in other animals [Jékely, 2013; Hentze et al., 2015; Christ et al., 2017]. However, octopus allatostatin lacks the critical LW motif diagnostic of this family of peptides (Figure S1), and this function may not be conserved. Our bioinformatics screen also predicted an IRELVamide peptide (Figure S1). This sequence resembles that of PRQFVamide and may represent an octopus variant.
